# Neurophysiology of perceptual decision-making and its alterations in attention-deficit hyperactivity disorder (ADHD)

**DOI:** 10.1101/2023.12.04.569762

**Authors:** Mana Biabani, Kevin Walsh, Shou-Han Zhou, Joseph Wagner, Alexandra Johnstone, Julia Paterson, Beth P. Johnson, Gerard M. Loughnane, Redmond G. O’Connell, Mark A. Bellgrove

## Abstract

Despite the prevalence of ADHD, efforts to develop a detailed understanding of the neuropsychology of this neurodevelopmental condition are complicated by the diversity of interindividual presentations and the inability of current clinical tests to distinguish between its sensory, attentional, arousal or motoric contributions. Identifying objective methods that can decompose the clinical heterogeneity of ADHD is a long-held goal that is hoped to advance our understanding of etiological processes and potentially aid the development of personalized treatment approaches. Here, we examine key neuropsychological components of ADHD within an electrophysiological (EEG) perceptual decision-making paradigm that is capable of isolating distinct neural signals of several key information processing stages necessary for sensory-guided actions from attentional selection to motor responses. We show that compared to typically developing children, children with ADHD displayed slower and less accurate performance, which was driven by the atypical dynamics of discrete electrophysiological signatures of attentional selection, the accumulation of sensory evidence, and strategic adjustments reflecting urgency of response. These findings offer an integrated account of decision-making in ADHD and establish discrete neural signals that can be used to understand the wide range of neuropsychological performance variations in individuals with ADHD.

**Significance Statement:** The efficacy of diagnostic and therapeutic pathways in ADHD is limited by our incomplete understanding of its neurological basis. One promising avenue of research is the search for basic neural mechanisms that may contribute to the variety of cognitive challenges associated with ADHD. We developed a mechanistic account of differences in a fundamental cognitive process by integrating across neurocognitive, neurophysiological (i.e., EEG), and computational levels of analysis. We detected distinct neural changes in ADHD that explained altered performance (e.g., slowed and less accurate responses). These included changes in neural patterns of attentional selection, sensory information processing, and response preparation. These findings enhance our understanding of the neurophysiological profile of ADHD and may offer potential targets for more effective, personalized interventions.

## Introduction

Attention-Deficit/Hyperactivity Disorder (ADHD) is a prevalent childhood-onset condition characterized by persistent inattentive, hyperactive and/or impulsive symptoms that significantly impact social relationships and quality of life (Barkley, 1997; Nigg, 2013; Sciberras et al., 2022). Early attempts to identify the neuropsychological characteristics of ADHD focused primarily on differences in higher-level cognitive processes associated with executive functioning such as response inhibition, working memory and cognitive flexibility (Pennington and Ozonoff, 1996; Barkley, 1997; Sergeant et al., 2002). However, ADHD is now recognised as a complex, heterogeneous, and multifactorial condition with contributions spanning multiple levels of processing (Sergeant et al., 2003; Willcutt et al., 2005; Coghill et al., 2014), including basic perceptual (Kim et al., 2014; Gonen-Yaacovi et al., 2016; Mihali et al., 2018; Panagiotidi et al., 2018) and neuromotor processes (Hurks et al., 2005; Rommelse et al., 2008; Kaiser et al., 2015; Goulardins et al., 2017). This raises the possibility that differences in higher-level processes may actually be the consequence of changes in more basic mechanisms that support downstream functions (Rommelse et al., 2007). Many of these component processes have been identified based on hallmark differences in performance on reaction time tasks, where ADHD participants typically exhibit reduced accuracy and slower, more variable reaction times (Bellgrove et al., 2005; Johnson et al., 2007; Karalunas and Huang-Pollock, 2013). However, since these behavioural outputs are the product of multiple processes (e.g., sensory encoding, evidence accumulation, motor preparation, urgency, decision bias and strategy), it is difficult to develop mechanistic accounts based on behavioural differences alone.

Sequential sampling models, like the Drift-Diffusion Model (DDM) (Ratcliff and McKoon, 2008), provide a powerful theoretical framework that can help to disentangle these influences by recovering latent psychological processes from their behavioural output (Forstmann et al., 2016). These models conceptualise decision-making as a dynamic process of the accumulation of noisy sensory evidence over time until a decision threshold is reached, and a response is initiated. Studies that have applied these models to the data of children with ADHD have identified slower accumulation of sensory evidence (reflected in a reduced drift rate parameter), no differences in bound adjustments (Mulder et al., 2010) and mixed evidence for differences in the non-decision time parameter which incorporates delays associated with stimulus encoding and motor execution (Huang-Pollock et al., 2012, 2017; Karalunas et al., 2012, 2014, 2018). Although these models highlight distinct decision-making mechanisms that are potentially altered in ADHD, they offer little insight into the underlying neurophysiological mechanisms.

Building on foundational work in monkey neurophysiology (Gold and Shadlen, 2007; Hanks and Summerfield, 2017), human research now capitalises on EEG paradigms to decompose simple decisions into their neural components, non-invasively mapping key information processing stages in decision-making (O’Connell and Kelly, 2021). These neural signals index processes representing candidate differences in decision-making in ADHD, including pre-target attentional engagement (alpha power) (Kelly and O’Connell, 2013), early attentional selection (N2c) (Loughnane et al., 2016), dynamic urgency (contingent negative variation; CNV) (Devine et al., 2019), and evidence accumulation (centroparietal positivity; CPP). As the neural marker of the evidence accumulation process, the behaviour of CPP is consistent with the predictions of sequential sampling models. Specifically, its build-up rate scales with evidence strength and predicts reaction time, while its peak amplitude, occurring at response, varies with prior knowledge and time pressure (Kelly and O’Connell, 2015; O’Connell et al., 2018).

Here, we sought to develop an integrated account of the neurophysiology of ADHD by linking these distinct EEG signals to mechanisms associated with performance of a perceptual decision-making task. We first aimed to establish linkages between EEG metrics of decision making with behaviour and DDM parameters in children with and without ADHD, using the Hierarchical DDM approach (Wiecki et al., 2013). Second, we aimed to characterise the dynamics of decision-making signals in ADHD, developing a mechanistic account that captures individual variations in performance. This comprehensive analysis allowed us to explore the neural, cognitive, and computational factors that govern decision-making in the context of ADHD.

## Materials and Methods

**Participants.** The study recruited a total of 79 right-handed individuals with normal or corrected-to-normal vision who were aged between 8 and 17 years, including 37 participants with ADHD (13 female; Mean_age_ = 13.45 years ± SD_age_ = 2.026_)_ and 42 healthy controls (18 female; Mean_age_ = 13.46 years ± SD_age_ = 1.93). Data were collected at the University of Queensland (UQ) (n = 58) and Monash University (n = 21) in Australia following identical experimental protocols. Ethical approval was obtained from the human research ethics committees of both universities, and the study was conducted in accordance with approved guidelines. For the ADHD group, inclusion criteria required previous diagnosis by a specialist (e.g., psychiatrist or paediatrician), confirmed using the Anxiety Disorders Interview Schedule for DSM-IV (A-DISC child version for UQ participants) or The Development and Well-being Assessment (DAWBA, for Monash participants) and the Global Index on the Conners’ Parent Rating Scale-Revised: Long Version (CPRS-R: L; T-Score > 65 from the mean). The participants with ADHD completed a 48-hour washout of ADHD-related medications before testing. Typically developing children were free of clinical diagnoses and had Conners’ Global Index T-Scores < 65. Informed consent from parents or guardians and assent was obtained from the participant prior to testing. **Figure 1** presents the clinical characteristics of the individuals in the two groups.

**Figure 1.**
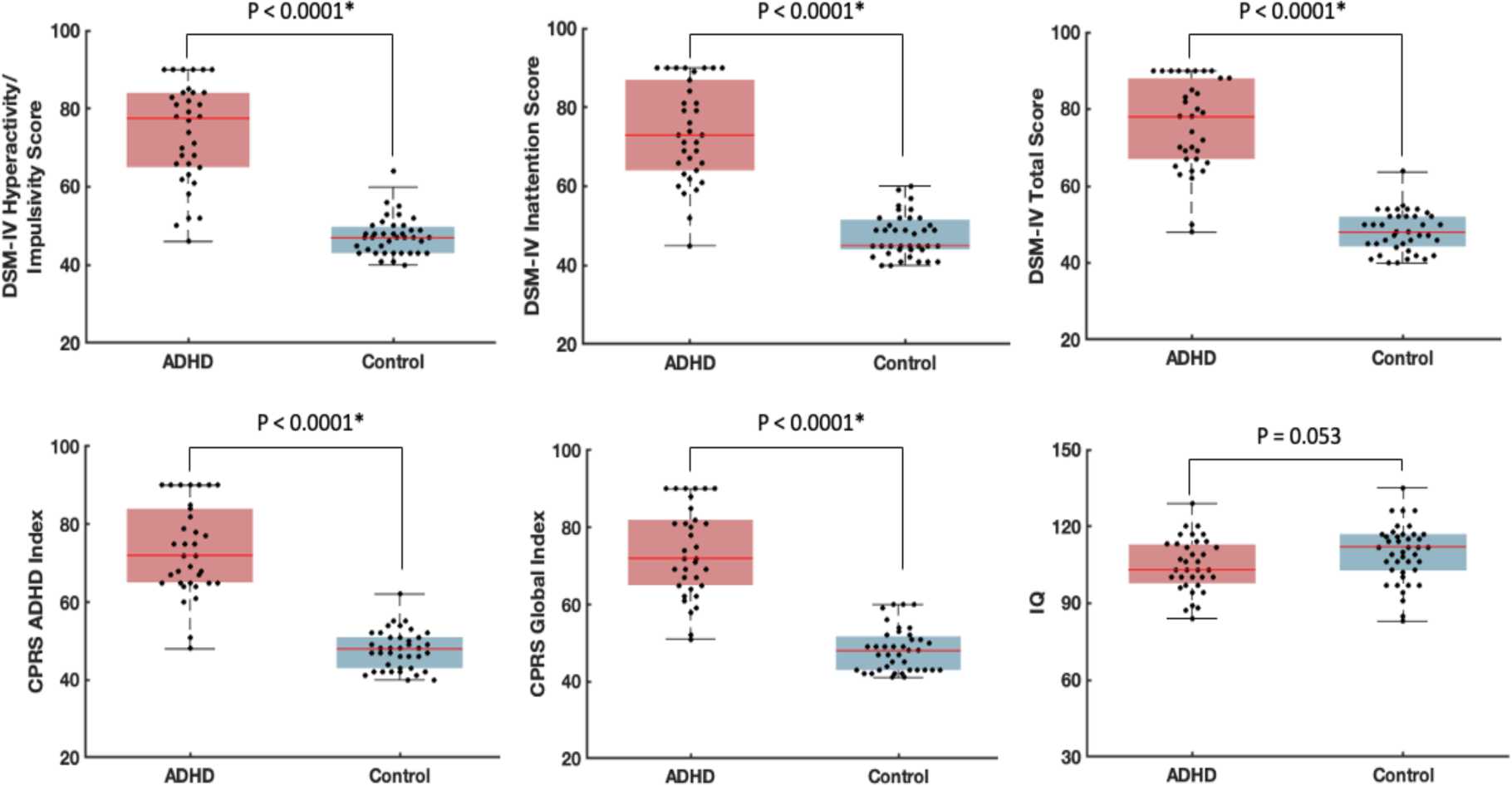
Clinical characteristics of participants in each group. P: P-Value from Wilcoxon-Signed Rank Test comparing the two groups. Every data point in the box and whisker plots corresponds to the clinical score for one individual. The shaded boxes indicate the range between the 25th and 75th percentiles of the scores, whereas the red horizontal lines inside the boxes represent the median score. * Indicates statistically significant differences between groups.

### Experimental Protocol

Participants were seated in a darkened sound-attenuated room positioned at a viewing distance of approximately 56 cm from a 21-inch CRT monitor (resolution: 1024 x 768, refresh rate: 85 Hz) and instructed to perform a bilateral random dot motion perceptual decision-making task. The task was run through MATLAB’s psychophysics toolbox extension on a 32-bit Windows XP computer (Brainard and Vision, 1997). A chin rest was employed to stabilise participants’ heads and maintain a constant visual angle throughout the task. An Eyelink 1000 eye tracking system (SR Research, Ottawa, ON, Canada) was used to monitor gaze at fixation.

In this task (Figure 2), participants fixated on a centrally presented 5×5-pixel square dot while simultaneously monitoring two circular patches, one per hemifield. These peripheral patches were eight degrees in diameter and contained 150 randomly moving 6×6-pixel dots. The centre of each patch was situated 4° below and 10° to the left or right of the central fixation dot to maintain an optimum visual angle for both hemifields. At pseudo-random intervals of either 3.06, 5.17 or 7.29 seconds, a subset of the dots transitioned to coherent downward motion for 1.88 seconds (Stefanac et al., 2021). These dots were randomly selected to move downwards by 0.282 degrees per frame (6 degrees per second). The coherent downward motion occurred once per trial and appeared with equal probability in either the left or right hemifield patch. Participants were instructed to respond promptly by pressing both mouse buttons simultaneously with their thumbs upon detecting the downward motion, employing a double thumb click. To capture robust accuracy and reaction time contrasts between groups, a relatively low coherence level of 30% was chosen for this task, deviating from previous studies using the same paradigm (Loughnane et al., 2016; Stefanac et al., 2021). Participants completed 200 trials of the task, divided into 10 blocks of 20 coherent motion trials each, with intermittent breaks to reduce fatigue.

**Figure 2.**
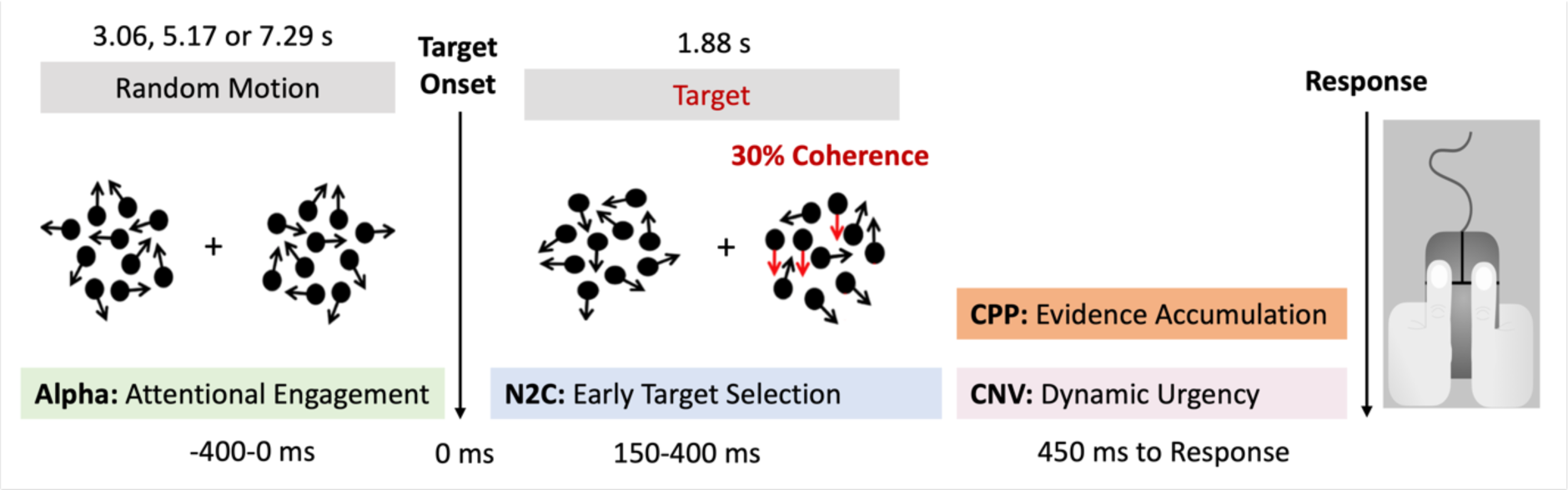
Depiction of the random-dot motion detection task and the obtained neural measures.

### EEG acquisition and pre-processing

Continuous EEG data were acquired from 64 scalp electrodes (10-10 layout) using BioSemi Active II (Neurospec, Switzerland) digitised at 1024 Hz (at the University of Queensland) or a BrainAmp DC system (BrainProducts, Germany) digitised at 500 Hz (at Monash University). Data were analysed using custom scripts in MATLAB (The MathWorks, Inc.) and EEGLAB toolbox (Delorme and Makeig, 2004). EEG recordings from the two locations were combined by down-sampling the data collected in Queensland to 500 Hz. Signals were low-pass filtered up to a 35 Hz cut-off using Hamming windowed-sinc FIR filter and no high pass filter was applied. Noisy channels were then interpolated using spherical spline interpolation and the data were re-referenced to the common average. Target epochs were extracted using a window of - 800 ms to 1880 ms around the onset of the target stimulus (coherent motion) and baseline corrected at -100 to 0 ms. The behavioural measures of reaction time (RT) and accuracy were extracted as the average time to respond to a target in milliseconds (ms) and the percentage of correctly identified targets, respectively. Trials were rejected if any of the following occurred: 1) the central gaze fixation was broken by blinking or vertical/horizontal eye movement greater than three degrees; 2) recordings from any electrode exceeded ±100 μV; 3) RTs were faster than 200 ms (pre-emptive responses) or slower than 1880 ms (responses after the offset of coherent motion). Missed targets were defined as either responses that took longer than 1880 ms or complete absence of responses. Hit rate was measured as the percentage of trials with valid responses.

ERPs for each individual were extracted from the average of single-trial epochs. For each individual, we isolated four distinct and previously validated EEG signatures of decision-making processes (Brosnan et al., 2020): pre-target attentional engagement (alpha power), early target selection (N2c peak latency and amplitude (Loughnane et al., 2016), evidence accumulation (CPP onset latency, slope, and amplitude (O’connell et al., 2012; McGovern et al., 2018; Steinemann et al., 2018), and dynamic urgency (CNV slope and amplitude (Devine et al., 2019). Pre-target alpha power was computed using the temporal spectral evolution approach, in which all epochs (-1000 ms to 2080 ms) were bandpass filtered at 8–13 Hz, rectified, and trimmed by 200 ms at both ends of the epoch (target epoch: -800:1880 ms) to eliminate filter warm-up artefacts. Subsequently, the data were smoothed by averaging within a 100-ms moving window shifting forward in 50-ms steps throughout each epoch. Mean alpha power was extracted bilaterally at peak electrodes - PO3/PO7 (left hemisphere) and PO4/PO8 (right hemisphere) - from 400 to 0 ms prior to the target onset and was baseline-corrected using the period of 700 to 400 ms before the target onset (Brosnan et al., 2020). For N2c components, peak negative amplitude was measured contralateral to the target hemifield, from electrodes P7 and P8 between 150 ms and 400 ms post target onset (Stefanac et al., 2021). CPP was extracted from the average of the potentials recorded at the peak electrodes - Pz and POz - and CNV was measured at FCz. The amplitude of each component was determined by calculating the mean signal amplitude within the same 100-ms window preceding the response. CPP onset latency was derived by performing point-by-point one-sample t-tests against zero over stimulus-locked trials for each individual. The onset was defined as the first point in time when the amplitude reached the significance level of 0.05 for 25 consecutive samples (Foxe and Simpson, 2002). The build-up rate of the CPP was measured as the slope of a straight line fitted to the response-locked signal at -450ms to -50ms(Loughnane et al., 2016; Zhou et al., 2021). The variation of the task used in this study, involving double thumb clicks, did not allow us to extract motoric signals, such as lateralized beta activity.

### Drift Diffusion Modelling

The drift diffusion model was fitted to the behaviour of all participants (both ADHD and typically developing children) at an individual level (Ratcliff et al., 2018). The response time data for each individual was first split into six equal speed bins defined by five quantiles (0.1, 0.3, 0.5, 0.7 and 0.9), resulting in four 20% bins and two 10% bins. Together with a single bin containing the number of missed responses, these seven bins were then used to fit the drift diffusion model using the G-square method of the hDDM package (Wiecki et al., 2013; Ratcliff et al., 2016; de Gee et al., 2020). This method is a variant of the chi-squared method and was chosen for its efficiency, the availability of significant trial data for each individual, its robustness to outliers and its success in previous similar experiments (Ratcliff et al., 2016; Myers et al., 2022). The G-square statistics is defined as:

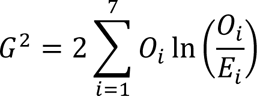

where *i* ∈ *N* represents the quantile number. The variable *O_i_* ∈ *R* represents the number of observations in each bin (in this case: 0.1, 0.2, 0.2, 0.2, 0.2 and 0.1 of the total number of observations), and *E_i_* ∈ *R* represents the expected number of observations in each bin, as predicted by the drift diffusion model. The expected number of observations is determined by first inserting the simulated response times into a drift diffusion model cumulative probability function to obtain the expected cumulative probability up to the five quantiles. Then, the proportion of simulated responses between each quantile is calculated by subtracting the cumulative probabilities for each successive quantile from the next highest quantile. This proportion is then multiplied by the total number of observations to obtain the expected frequencies. The drift-diffusion model parameters *a*, *v* and *t* were determined by minimising the G-square statistic using the modified Powell method (Powell, 1964). The fitted DDM assumed that the response caution/urgency, mean drift rate across trials, and non-decision time varied with the response bins. Further details can be found in (Ratcliff et al., 2018).

### Statistical Analysis

Significant outliers were winsorised to the 5th percentile (for the lower outliers) and the 95th percentile (for the upper outliers) in each participant group to improve normality of distributions. CPP onset could not be detected for five individuals in each group due to noisy signals, so the missing values were replaced with the group median.

First, we adopted a hierarchical regression approach to investigate the association between the EEG components and both behavioural and DDM outcomes. Separate hierarchical linear regression models were applied for each behavioural and DDM measure as a function of the EEG signals. EEG components were sequentially entered into the regression models, following the temporal order of the perceptual decision-making processes: pre-target attentional engagement (alpha power), early target selection (N2c peak latency and amplitude), evidence accumulation (CPP onset, slope and amplitude) and dynamic urgency (CNV slope and amplitude). This hierarchical entry method allowed us to evaluate whether each individual neurophysiological signal contributed to the model fit for behavioural performance or DDM parameters beyond the preceding signals in the temporal sequence. Any neurophysiological signals that improved the model fit were incorporated into a separate regression model to obtain accurate parameter estimates. Next, Spearman’s partial correlation analyses were employed, controlling for the effects of group (ADHD, control) and recruitment site (Monash, UQ), to evaluate the magnitude and orientation of the relationship between EEG components and both behavioural outcomes and DDM parameters.

To determine any differences in decision-making processes in ADHD versus typically developing children, a multivariate analysis of covariance (MANCOVA) was conducted to compare the two groups on behavioural measures (RT, Hit Rate and Miss Rate), EEG signatures including alpha (power), N2c (peak latency, amplitude), CPP (onset, amplitude, slope) and CNV (amplitude, slope), and the DDM parameters (drift rate, decision threshold, non-decision time), while controlling for the effect of recruitment site. We also examined whether the EEG signals were related to ADHD symptom scores while accounting for the effects of group and site. For this analysis we employed five separate linear regression models, followed by false discovery rate (FDR) adjustment, each with one ADHD symptom domain as the dependent variable (i.e., DSM-IV hyperactivity/impulsivity score, DSM-IV inattention score, DSM-IV total score, CPRS ADHD Index, CPRS Global Index), while EEG signatures were entered into the model hierarchically.

## Results

We first examined the relationship between each EEG signal (alpha, N2c, CPP and CNV) and variations in behaviour using a hierarchical regression model. In the initial step of the model, group and site were entered as nuisance variables. In the subsequent steps, we sequentially incorporated the neural markers of attentional engagement (alpha power), target selection (N2c: peak latency and amplitude), evidence accumulation (CPP: onset, amplitude, and slope), and dynamic urgency (CNV: amplitude and slope) into the model. This hierarchical methodology allowed us to control for the chronological order of neural processes in perceptual decision-making and examine the incremental predictive power of different signals. Although the existing literature suggests connections between certain EEG signals and behavioural/DDM measures (Yau et al., 2021), we avoided making *a-priori* assumptions about these relationships in our analysis. Instead, we systematically examined each component to evaluate their impact and uncover any latent patterns and interactions.

### Neural metrics of sensory evidence accumulation capture variations in reaction time

The neural signatures of the decision process collectively accounted for a substantial 46% of the variance in RT. Adding each of the neural components resulted in a significant improvement in the model fit (alpha power: R^2^_adj_ = 0.16, F (3,75) = 5.87, p = 0.001; N2c latency: R^2^_adj_ = 0.16, F (4,74) = 4.63, p = 0.002; N2c amplitude: R^2^_adj_ = 0.15, F (5,73) = 3.77, p = 0.004; CPP onset: R^2^_adj_ = 0.28, F (6,72) = 5.98, p <0.001; CPP slope: R^2^_adj_ = 0.37, F (7,71) = 7.59, p <0.001; CPP amplitude: R^2^_adj_ = 0.47, F (8,70) = 9.53, p <0.001; CNV slope: R^2^_adj_ = 0.47, F (9,69) = 8.56, p <0.001; CNV amplitude: R^2^_adj_ = 0.46, F (10,68) = 7.63, p <0.001). The analysis of coefficients revealed that alpha power (stand. β = 0.18, t = 2.07, p = 0.04) and all CPP components (onset: stand. β = 0.47, t = 4.81, p < 0.001; slope: stand. β = -0.84, t = -4.37, p < 0.001; amplitude: stand. β = 0.61, t = 2.87, p = 0.005) had independent predictive power for RT, highlighting their potential as robust markers of changes in decision making processes.

### Neural metrics of sensory evidence accumulation capture variations in Miss Rate

The second model examined the neural predictors of Miss Rate and yielded comparable outcomes to those of RT. The EEG signals collectively accounted for 26% of variations in Miss Rate. All neural components significantly improved model fit (alpha power: R^2^_adj_ = 0.15, F (3,75) = 5.47, p = 0.002; N2c latency: R^2^_adj_ = 0.14, F (4,74) = 4.09, p = 0.005; N2c amplitude: R^2^_adj_ = 0.14, F (5,73) = 3.52, p = 0.007; CPP onset: R^2^_adj_ = 0.26, F (6,72) = 5.50, p < 0.001; CPP slope: R^2^_adj_ = 0.25, F (7,71) = 4.71, p < 0.001; CPP amplitude: R^2^_adj_ = 0.24, F (8,70) = 4.09, p < 0.001; CNV slope: R^2^_adj_ = 0.24, F (9,69) = 3.76, p < 0.001; CNV amplitude: R^2^_adj_ = 0.27, F (10,68) = 3.76, p < 0.001) but only CPP onset demonstrated independent predictive power for Miss Rate (stand. β = 0.45, t = 3.99, p < 0.001), underlying its significance in determining performance on this particular task.

### Neural metrics of sensory evidence accumulation capture variations in Hit Rate

In the third model we explored the neural predictors of Hit Rate. The examined EEG metrics collectively accounted for 26% of the variance in Hit Rate. In line with the results from RT and Miss Rate, adding each metric substantially enhanced the model’s overall fit (alpha power: R^2^_adj_ = 0.14, F (3,75) = 5.18, p = 0.003; N2c latency: R^2^_adj_ = 0.13, F (4,74) = 3.85, p = 0.007; N2c amplitude: R^2^_adj_ = 0.13, F (5,73) = 3.31, p = 0.009; CPP onset: R^2^_adj_ = 0.23, F (6,72) = 4.84, p < 0.001; CPP slope: R^2^_adj_ = 0.22, F (7,71) = 4.16, p < 0.001; CPP amplitude: R^2^_adj_ = 0.22, F (8,70) = 3.67, p = 0.001; CNV slope: R^2^_adj_ = 0.21, F (9,69) = 3.31, p = 0.002; CNV amplitude: R^2^_adj_ = 0.26, F (10,68) = 3.67, p < 0.001), and CPP onset (stand. β = -0.45, t = -3.93, p < 0.001) and CNV amplitude (stand. β = -0.34, t = -2.27, p = 0.03) made significant individual contributions to the prediction of Hit Rate.

Together, these results provide compelling evidence that the EEG metrics of target selection (N2c), evidence accumulation (CPP), and dynamic urgency (CNV) collectively exhibit predictive power for decision-making performance across the three behavioural measures (RT, Miss Rate, Hit Rate). This underscores the importance of considering multiple neural components when investigating and interpreting decision-making processes. Although the neural signals showed varying contributions to each behavioural measure, CPP onset emerged as an independent predictor for performance variations across all the three measures. Figure 3 illustrates the relationships between CPP onset and performance, along with the Pearson’s correlation coefficients.

**Figure 3.**
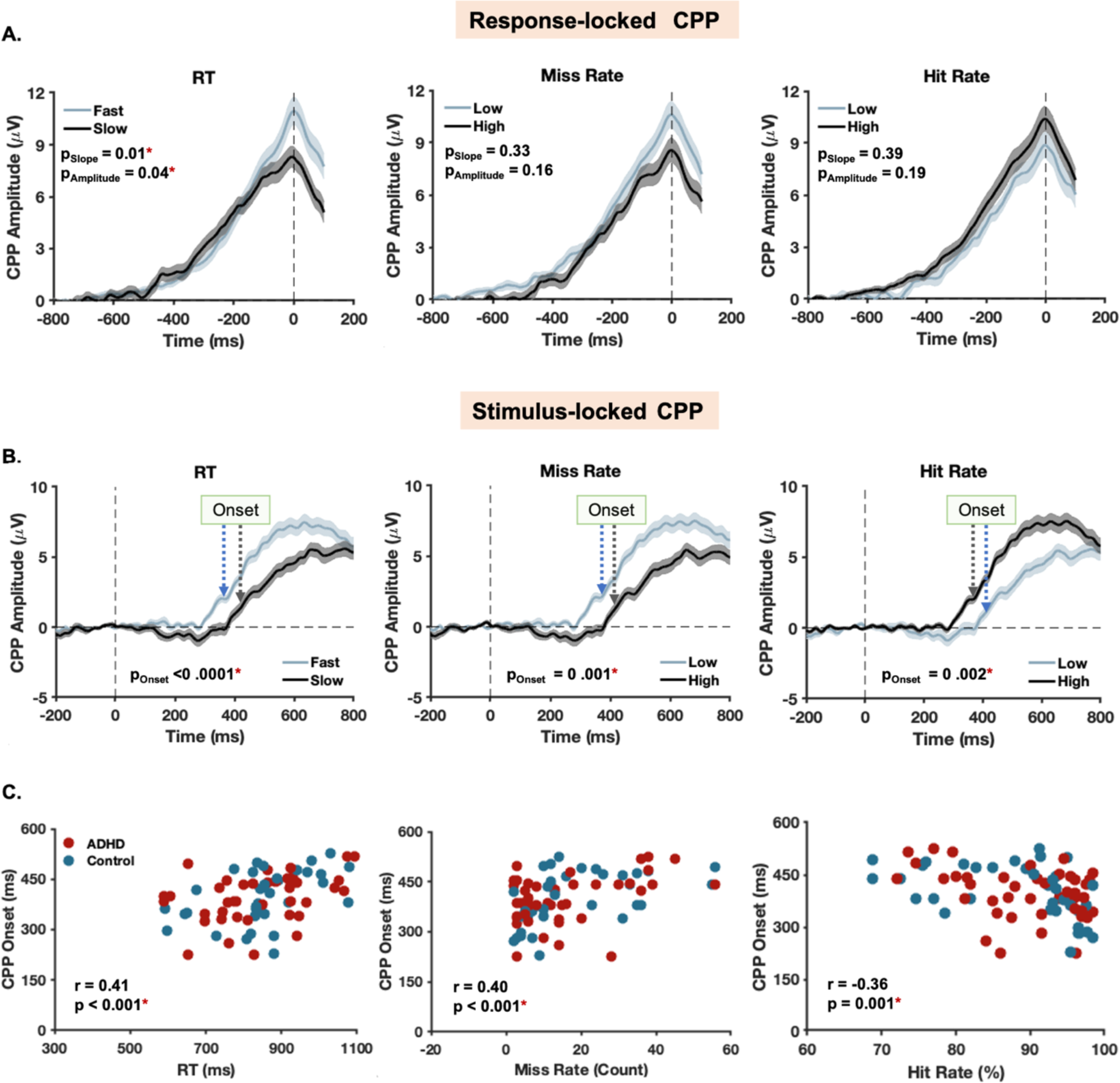
Relationship between CPP dynamics and performance. **A-B.** Differences in CPP signal between individuals with different levels of performance. CPP amplitude and slope are measured from response-locked CPP (A) and CPP onset is from stimulus-locked CPP (B). Participants are binned by the behavioural measure using a median split of the data. The thick line in the graph is the group-averaged waveform and the shaded areas represent changes in the standard error of mean over time. The vertical dashed lines marked *onset* compare the average onsets between the groups. P: p-value from Wilcoxon-Signed Rank Test comparing the two groups. * indicate statistical significance (P<0.05) in group difference. **C.** The relationship between CPP onset (derived from stimulus-locked CPP), which demonstrated significant predictive power for all behavioural measures, and performance. P and r: p-value and coefficient from partial Pearson’s correlation analysis while controlling for group and site.

### Collective predictive power of neural metrics on DDM parameters

Next, we examined whether the EEG signatures of decision-making processes were associated with DDM parameters fitted to the behavioural measures by modelling each DDM parameter as a function of the EEG signals in hierarchical regression analysis. Group and site were entered as nuisance factors at the first stage of each model.

### Non-decision Time (t)

All of the neural metrics of decision-making significantly improved the model fit for the non-decision time parameter of the DDM (Figure 4A) explaining 24% of the variance (N2c latency: R^2^_adj_ = 0.24, F (3,75) = 9.11, p < 0.001; N2c amplitude: R^2^_adj_ = 0.24, F (4,74) = 7.10, p < 0.001; CPP onset: R^2^_adj_ = 0.24, F (5,73) = 5.97, p < 0.001; CPP slope: R^2^_adj_ = 0.24, F (6,72) = 5.03, p < 0.001; CPP amplitude: R^2^_adj_ = 0.23, F (7,71) = 4.37, p < 0.001; CNV slope: R^2^_adj_ = 0.23, F (8,70) = 3.80, p < 0.001; CNV amplitude: R^2^_adj_ = 0.24, F (9,69) = 3.73, p < 0.001). However, none of the EEG signals demonstrated independent predictive power for this DDM parameter (all p > 0.05).

**Figure 4.**
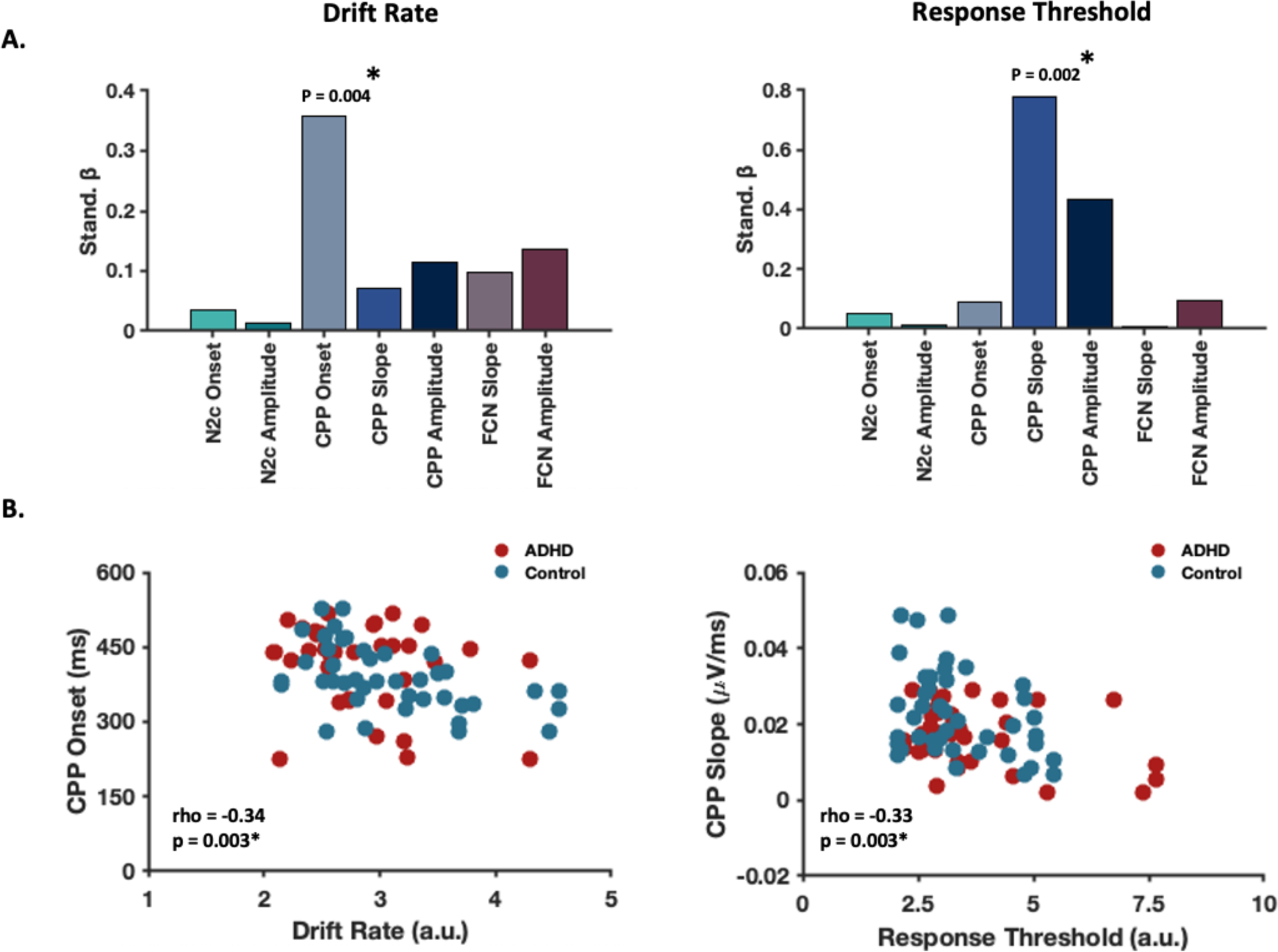
Relationships between DDM parameters and EEG signals. **A.** Results from the hierarchical regression models of DDM parameters indicating the predictive power of the EEG signals. * indicates that the EEG signal indicated independent predictive power for the DDM measure. Note that the standardised beta values are displayed as absolute values for the purpose of visualisation and the plots do not include the nuisance variables (group and site) that were entered into the model in the first step. **B.** Results from partial Spearman’s correlation analysis depicting the relationship between the CPP component and DDM measures.

### Drift Rate (ν)

Similarly, the model fit for the drift rate parameter was significantly improved when each of the EEG metrics were added into the model (Figure 4A). This model explained 15% of the variance in drift rate (N2c latency: R^2^_adj_ = 0.09, F (3,75) = 9.11, p = 0.02; N2c amplitude: R^2^_adj_ = 0.08, F (4,74) = 7.10, p = 0.04; CPP onset: R^2^_adj_ = 0.17, F (5,73) = 5.97, p = 0.002; CPP slope: R^2^_adj_ = 0.18, F (6,72) = 5.03, p = 0.002; CPP amplitude: R^2^_adj_ = 0.17, F (7,71) = 4.37, p = 0.005; CNV slope: R^2^_adj_ = 0.16, F (8,70) = 3.80, p = 0.01; CNV amplitude: R^2^_adj_ = 0.15, F (9,69) = 3.73, p < 0.01) and CPP onset accounted for independent variation in drift rate (stand. β = -0.36, t = -2.95, p = 0.004).

### Response Threshold (a)

The model fit for the response threshold parameter was significantly improved by the inclusion of CPP slope (R^2^_adj_ = 0.14, F (6,72) = 3.70, p = 0.008) and amplitude (R^2^_adj_ = 0.19, F (7,71) = 3.22, p = 0.002) and CNV dynamics (slope: R^2^_adj_ = 0.19, F (8,70) = 3.22, p = 0.004; amplitude: R^2^_adj_ = 0.18, F (9,69) = 2.87, p = 0.06), but not CPP onset (R^2^_adj_ = 0.02, F (5,73) = 1.38, p = 0.24) or N2c measures (latency: R^2^_adj_ = 0.05, F (3,75) = 2.4, p = 0.08; amplitude: R^2^_adj_ = 0.04, F(4,74) = 1.75, p = 0.15) (Figure 4A). To obtain accurate parameter estimates for the association between response threshold and neural components, without the influence of the non-informative signals, CPP (slope and amplitude) and CNV measures were entered into a separate model with the nuisance factors included. This model explained 20% of the variation and CPP slope showed independent predictive power for response threshold (stand. β = -0.12, t = -3.30, p = 0.002).

The results above collectively provide evidence supporting associations between neural metrics of decision-making as measured with EEG, and DDM parameters derived from behavioural outcomes. Among all EEG components, the dynamics of CPP emerged as the strongest contributor to the variations in DDM parameters. Figure 4B illustrates the relationship between CPP dynamics and DDM parameters employing partial Spearman’s correlation analysis, for non-normally distributed data, while controlling for group and site. Notably, a significant correlation was found between CPP onset and drift rate, as well as between CPP slope and response threshold.

### Neurobehavioural characteristics of decision-making in ADHD

To investigate the distinctions in behavioural, DDM and electrophysiological measures of decision-making between the ADHD and control groups, we conducted MANCOVA with site as a covariate. As the Box’s M test score was significant (p = 0.04), we used Pillai’s trace statistic, which is considered to be most robust against type I error in MANCOVA (Olson, 1976; Scheiner, 2020). The results revealed significant main effects of both Group (Pillai’s Trace = 0.31, F (14, 63) = 2.05, p = 0.02, partial η2 = 0.31) and Site (Pillai’s Trace = 0.46, F (14, 63) = 3.90, p <0.001, partial η2 = 0.46).

Table 1 summarises pairwise comparisons of all the measures between the ADHD and control groups. Significant differences were observed between the two groups in all measures of performance and EEG (in at least one feature of the signal), with the exception of pre-target alpha. Figure 5 illustrates the differences in spatiotemporal patterns of EEG signals between the two groups. Despite individuals with ADHD showing a notable reduction in the DDM drift rate parameter in the expected direction, neither it nor the other DDM parameter estimates showed significant differences between the two groups.

**Table 1.** Pairwise comparisons of measures between the ADHD and typically developing groups.

| Measures | Mean Difference | Std. Error | Sig. | 95% Confidence Interval |  |
| --- | --- | --- | --- | --- | --- |
| **RT (ms)** | 58.57 | 28.45 | **0.04\*** | 1.91 | 115.24 |
| **Hit Rate (%)** | -3.70 | 1.73 | **0.03\*** | -7.14 | -0.26 |
| **Miss Rate (count)** | 7.70 | 2.91 | **0.01\*** | 1.90 | 13.51 |
| Drift Rate (a.u.) | -0.24 | 0.13 | 0.07 | -0.50 | 0.03 |
| Non-Decision Time (ms) | 0.26 | 0.23 | 0.38 | -0.19 | 0.72 |
| Response Threshold | 0.33 | 0.28 | 0.24 | -0.23 | 0.89 |
| Pre-target alpha †($\mu V^2$) | -0.36 | 0.05 | 0.51 | -0.14 | 0.72 |
| **N2c Latency (ms)** | -24.93 | 10.95 | **0.02\*** | -46.75 | -3.13 |
| **N2c Amplitude ($\mu V$)** | 0.75 | 0.32 | **0.02\*** | 0.11 | 1.40 |
| CPP Onset (ms) | 25.50 | 16.55 | 0.13 | -7.46 | 58.45 |
| **CPP Slope ($\mu V/ms$)** | -0.01 | 0.002 | **0.02\*** | -0.01 | -0.001 |
| **CPP Amplitude ($\mu V$)** | -2.36 | 0.88 | **0.009\*** | -4.12 | -0.61 |
| CNV Amplitude ($\mu V$) | 1.29 | 1.04 | 0.21 | -0.78 | 3.37 |
| **CNV Slope ($\mu V/ms$)** | 0.02 | 0.01 | **0.03\*** | 0.002 | 0.04 |

\* The mean difference is significant at an alpha level of 0.05, which survived following FDR correction for multiple comparisons in each category of measures (behaviour, EEG and DDM). † Mean values are multiplied by $10^{15}$ .

**Figure 5.**
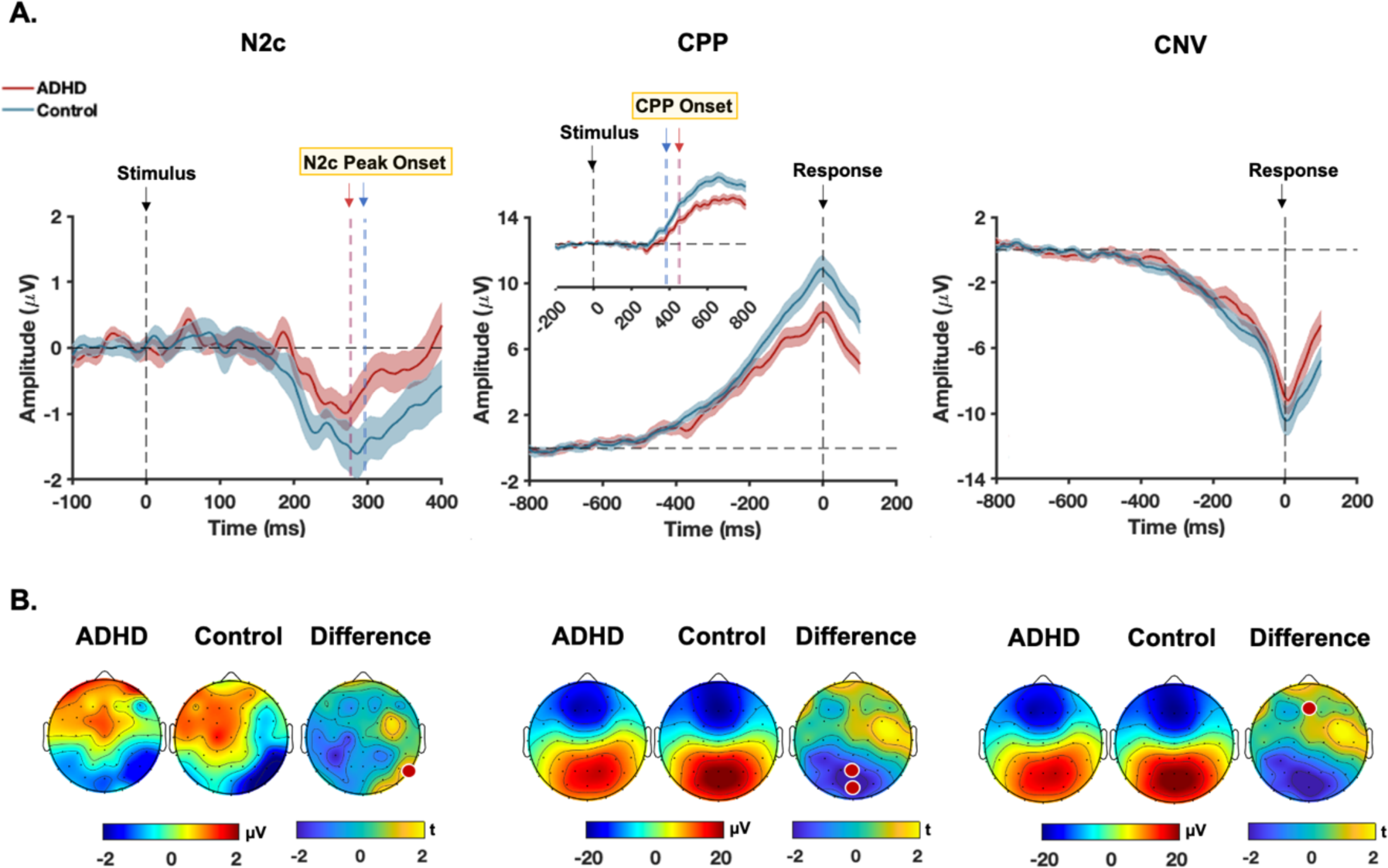
EEG signals of decision making in ADHD and typically developing groups. **A.** Group average EEG signal waveforms for each neural signature of the decision process. The middle graph displays the response-locked CPP, from which we derived measurements of CPP amplitude and slope. The inset graph depicts the stimulus-locked CPP, used to determine CPP onset. The thick line in the graphs is the group-averaged waveform and the shaded areas represent the standard error of mean at each point of time. The vertical dashed line at the zero point indicates the onset of the target stimulus for N2c and stimulus-locked CPP and represents the response time for response-locked CPP and CNV. The horizontal dashed line represents the baseline level of EEG activity. The vertical dashed lines marked *onset* compare the average onsets between the groups. **B.** The scalp maps depict the potential distribution for each group at the time of peak amplitude, and difference maps demonstrate the distribution of t-values resulting from t-tests comparing the two groups. The electrodes highlighted in red indicate the specific electrodes used for the line graphs and subsequent analysis. The scalp maps for CPP represent response-locked signals used to measure amplitude. Statistical outcomes are detailed in Table 1.

### Group differences in reaction time are mediated by variations in evidence accumulation

Given the predictive power of the CPP for decision-making performance and its capacity to distinguish between groups, we tested whether CPP dynamics mediated the performance differences observed in ADHD. Because the mediation effects of the different CPP metrics were likely related, for each behavioural measure, we jointly tested all the three mediators in one model to assess simultaneous effects more accurately (MacKinnon et al., 2000, 2007). Bootstrapped mediation analyses with 5000 samples (bias-corrected percentile; with site as confounding factor) revealed that the inter-subject variation in RT for the ADHD group, at least in part, depends on individual differences in the efficiency of evidence accumulation (Table 2).

**Table 2.**
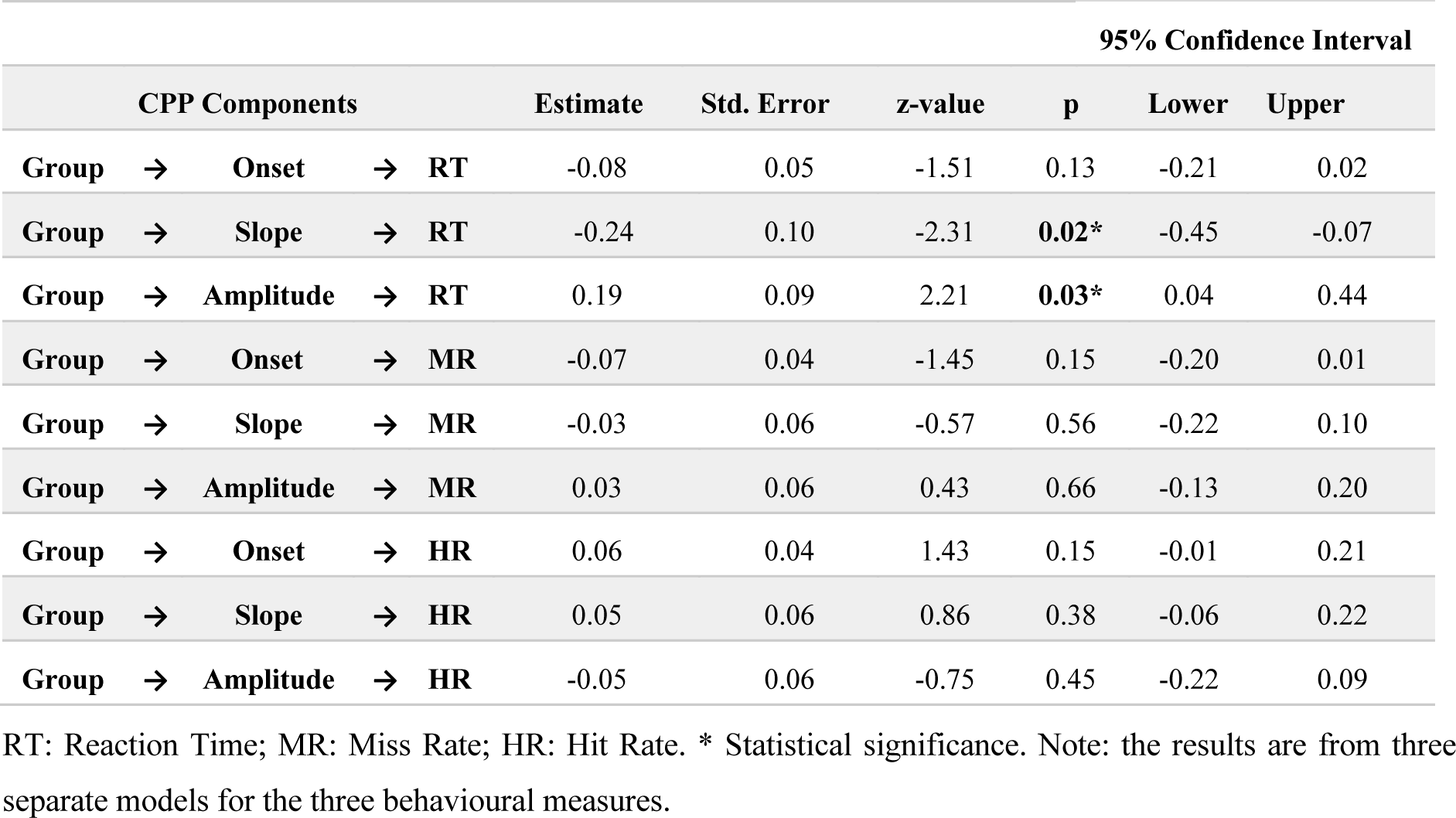
Mediation of CPP components on the impact of group on behaviour.

### Clinical scores

Finally, we sought to determine whether the EEG signatures could serve as predictors for the clinical scores. Hierarchical regression models revealed that variations in the EEG signatures collectively accounted for a significant portion of ∼70-80% (R^2^_adj_) variance in each clinical score. The model fit for each score was significantly improved by adding each neural metric (all p _FDR-corrected_<0.001). However, none of the EEG signals emerged as independent predictors for the clinical scores suggesting that these scores may reflect the interplay of multiple processing stages.

## Discussion

In this study, we aimed to develop a mechanistic account of ADHD-related changes in a fundamental cognitive process by integrating neurocognitive, neurophysiological, and computational levels of analysis, and identified distinct phenotypic signatures. First, our findings confirmed the link between performance and the EEG signatures of cognitive processes during perceptual decision-making, highlighting CPP dynamics (onset and slope) as robust, independent neural predictors across various measures (RT, miss and hit rates). Also, consistent with the literature (Rommelse et al., 2007; Karalunas and Huang-Pollock, 2013), the ADHD cohort demonstrated significantly slower RTs, a higher number of missed targets, and a reduced hit rate. In addition, children with ADHD exhibited altered dynamics in several neurophysiological signatures of the decision process including target selection (early and attenuated N2c), evidence accumulation (reduced CPP slope and amplitude) and anticipation of voluntary action (reduced CNV slope). Critically, the interplay of these neural signals explained meaningful inter-individual variation in performance and clinical outcomes. DDM parameters, however, demonstrated limited sensitivity in distinguishing ADHD from typically developing children, although significantly correlated with the CPP dynamics.

The N2c component is a key signature of early target selection mechanisms that support the decision process by facilitating enhanced processing of target features (Loughnane et al., 2016). Although the marginally earlier latency of the N2c observed in children with ADHD could be interpreted as enhanced target detection, its diminished amplitude coupled with the poorer performance of the ADHD group, more likely indicate premature processing of sensory information and/or changes in allocation of attentional resources. The N2c is closely related to the N2pc component elicited during visual search tasks, reflecting attentional impairment in ADHD, as indicated by altered timing and reduced amplitude (Cross-Villasana et al., 2015; Wang et al., 2016; Luo et al., 2019). Like the N2pc, the N2c functions as a general target selection signal emerging irrespective of the presence of distractors or the degree of spatiotemporal uncertainty (Loughnane et al., 2016). Although the N2c did not seem to act as an independent predictor of performance or clinical characteristics in ADHD, its significant contribution to models predicting both behavioural outcomes and DDM parameters suggests a partial link between ADHD-related decision-making impairments and the target selection mechanisms that support the decision process. N2c dynamics are relatively unexplored in the ADHD literature, but, given the centrality of attentional differences to ADHD, it merits further investigation.

There is evidence that a key mechanism that changes in ADHD may be the rate of basic information processing (Rommelse et al., 2007; Salum et al., 2014b, 2014a; Mihali et al., 2018). Indeed, efforts to model a variety of behavioural impairments associated with ADHD have consistently highlighted the rate of evidence accumulation as the core contributor to performance changes (Karalunas et al., 2012, 2014; Huang-Pollock et al., 2017, 2020; Weigard and Sripada, 2021). Our results complement this work by tracing the dynamics of an established neurophysiological index of evidence accumulation, the CPP. CPP dynamics not only exhibited robust predictive power for all three measures of cognitive performance, but also differed significantly between ADHD and typically developing individuals. In line with claims of inefficient evidence accumulation in ADHD, we found a reduced CPP slope and a smaller CPP amplitude in the ADHD group. Consistent with previous research showing that steeper CPP slopes predict faster reaction times and CPP amplitude is reliably greater for hits than for misses (O’connell et al., 2012; Kelly and O’Connell, 2013, 2015), this result provides neurophysiological evidence to support the hypothesis that suboptimal evidence accumulation or poorer signal-to-noise ratios in the processing of task-relevant information may contribute to differences in decision-making in ADHD.

To our knowledge, this study is the first to investigate CPP dynamics in ADHD. However, the P300 event-related potential^1^(Twomey et al., 2015; O’Connell and Kelly, 2021), also associated with evidence accumulation, has consistently been reported to have reduced amplitude in ADHD across a variety of tasks (Itagaki et al., 2011; Hasler et al., 2016; Kaiser et al., 2020). In fact, these effects are sufficiently robust that P300 dynamics have been proposed as potential ADHD biomarkers (Kaiser et al., 2020) and metrics for research on pharmacological treatment (Ogrim et al., 2016; Yamamuro et al., 2016; Peisch et al., 2021). Although the interpretation of these P300 effects varies across tasks, these results can be broadly characterised as reflecting suboptimal processing of task-relevant information. The CPP is thought to be functionally equivalent to the P300 (Itagaki et al., 2011), but studying the CPP offers several critical advantages over this previous work. Unlike the stimulus-locked P300, the slope and amplitude of CPP are estimated from the response-locked potentials accounting for the fact that the signal peaks at the time of response. Furthermore, the P300 analysis often overlooks the onset and build-up rate of the signal which are critical for understanding the neural processes underlying evidence accumulation. The present findings also align with research that indicates methylphenidate (MPH) enhances cognitive task performance by improving evidence accumulation. MPH is the mainstay treatment for ADHD and has been shown to normalise the reduced DDM drift rate in ADHD (Fosco et al., 2017). It also realigns P300 dynamics in neurocognitive (Peisch et al., 2021) and perceptual decision-making tasks (Loughnane et al., 2019). Additionally, preliminary evidence suggests that MPH enhances CPP slope in human EEG (Loughnane et al., 2019). The neural mechanisms by which MPH might enhance evidence accumulation are still largely unknown although some evidence from behavioural modelling studies suggests that it may regulate the suboptimal neural signal-to-noise ratios in children with ADHD (Ratcliff et al., 2009; Loughnane et al., 2019; Pertermann et al., 2019), suggesting that an increase in neural gain may account for effects observed on the P300. Future studies may yield a deeper understanding of the pharmacology of discrete processing stages underlying human choice behaviour by integrating the EEG paradigms and computational modelling approach employed in the present study with pharmacological manipulation.

Our computational modelling suggests that none of the drift diffusion parameters statistically differentiate the groups, despite a trend towards a reduced drift rate for the ADHD group. This stands in contrast to a number of reports in the literature employing a range of neurocognitive paradigms (Weigard and Sripada, 2021). One possibility is that the more basic perceptual decision-making task used in this study was less sensitive to drift rate differences than more cognitive demanding tasks (e.g., go/no-go, or continuous performance task). However, it should be noted that other studies with similar numbers of participants and trials, requiring simple decisions about basic stimuli (e.g. arrows) have reported drift rate effects associated with ADHD (Metin et al., 2013). It is worth noting that these studies have made specific assumptions regarding the drift diffusion model parameters, such as fixing the boundary condition, while the paradigm here makes no such assumption. It is therefore difficult to concretely attribute the absence of a drift rate effect to a particular aspect of the paradigm. Indeed, in the present study, multiple neural signals predicted changes of each DDM parameter suggesting that these parameters are the product of multiple decision processes.

Finally, our data revealed ADHD-related changes in CNV dynamics, which also contributed to the variation in behavioural performance. The CNV signal is commonly observed in target detection and choice response time tasks, which is associated with temporal preparation for anticipated events or volitional movements (Brunia and Van Boxtel, 2001; Van Rijn et al., 2011; Baker et al., 2012). This signal is influenced by dopaminergic systems (Birbaumer et al., 1990) and its attenuation has been widely reported in children (Banaschewski et al., 2003; Doehnert et al., 2013; Kaiser et al., 2020) and adults with ADHD (McLoughlin et al., 2010, 2011; Hasler et al., 2016). Indeed, this signal has been suggested as a robust neurophysiological marker of ADHD which effectively captures the underlying deficits in their preparatory motor processes (Doehnert et al., 2013; Kaiser et al., 2020). In the context of perceptual decision-making, the CNV is also described as a neural index of urgency which grows in a time-dependent but evidence-independent manner reflecting speed pressure in response (Devine et al., 2019). Given the slowed evidence accumulation in the ADHD group, one might expect an increased urgency to reach decision commitments as a compensatory mechanism. It appears that such strategic adjustment was not adaptive here as the ADHD group demonstrated poorer performance on average. Although numerous studies suggest that decision boundaries do not differ between ADHD and control groups, there is some evidence that they may have difficulty flexibly adjusting the inherent speed/accuracy trade-off in response to task demands (Mulder et al., 2010). It is possible that dysregulation of the timing mechanism associated with the CNV may contribute to this relative inflexibility.

The present study provides novel neurophysiological evidence linking differences in decision-making in ADHD to alterations in the dynamics and interplay of the neural signals indexing three key cognitive processes: target selection, decision formation and dynamic urgency. The results offer an integrated account of these changes and establish neural signals that can serve as critical guidance in constructing or constraining mechanistic accounts in future ADHD research.

## Acknowledgements

This work is supported by grants from the National Health and Medical Research Council (NHMRC) of Australia to M.A.B (#s2010899; 2025415)

## Footnotes

1 The CPP is typically observed during extended perceptual discrimination, while the P300 is evoked by discrete sensory events (e.g., an oddball stimulus).

